# Individualized Alpha-tACS for Modulating Pain Perception and Neural Oscillations: A Sham-Controlled Study in Healthy Participants

**DOI:** 10.1101/2025.01.16.633370

**Authors:** Yaser Fathi, Françoise Dissassuca, Gloria Ricci, Giulia Liberati

## Abstract

Pain encompasses sensory, affective, and cognitive dimensions, with neural oscillations increasingly recognized as key mechanisms in their integration. However, the underlying processes remain inadequately understood. Transcranial alternating current stimulation (tACS) offers a promising tool for modulating these oscillations, yet the widespread reliance on ‘one-size-fits-all’ tACS protocols with fixed frequencies has led to limited and contradictory findings on its efficacy in pain treatment. In this study, we employed individualized tACS at individual peak alpha frequency (IAF) over the primary motor cortex (M1) contralateral to the dominant arm of 38 healthy participants, in a within-subject, sham-controlled design, to investigate its effects on pain perception and neural oscillations. Sustained and periodic 0.2 Hz thermonociceptive stimuli were applied to the dominant forearm before and after tACS. We measured participants pain perception and heat pain thresholds (HPT) before and after tACS stimulation. Scalp electroencephalography (EEG) measurements were used to measure neural activity during thermonociceptive stimuli. To calculate IAF, we used a discriminative approach based on independent component analysis (ICA) to separate sensorimotor related IAF (SM-IAF). The results revealed an overall increase in pain perception and a decrease in HPT in both sham and active conditions, with no significant interactions between conditions. However, a trend toward reduced sensitization post-tACS was observed. Exploratory analyses indicated a significant tACS effect on HPT in women. Furthermore, a significant correlation was found between SM-IAF and HPT. These findings provide a novel perspective on advancing individualized neuromodulation approaches for pain and neurobiological disorders.

## 1. Introduction

Despite being a pervasive experience in everyday life, the exact mechanisms through which pain arises from human brain activity have not yet been unraveled. Pain involves the integration of sensory, affective, and cognitive dimensions, and growing evidence suggests that neural oscillations may serve as mechanisms facilitating this integration. Previous studies have highlighted that altered neural oscillations could be related to pain perception [14,17,25,28,34,35,39].

Sustained periodic thermonociceptive stimuli perceived as painful have been shown to selectively modulate oscillatory activity in distinct frequency bands, as revealed through both scalp electroencephalography (EEG) [8] and intracerebral electroencephalography (iEEG) [27]. Changes in alpha oscillations, specifically an increase in low-alpha activity, have been consistently reported in those suffering from chronic pain [14,43,49,50]. The relation between individual peak frequency of alpha oscillations (IAF), and experimental pain perception has been investigated in several studies [7,11–13,30,33,47]. Furman et al. found that sensorimotor IAF (SM-IAF) is a reliable biomarker of prolonged pain sensitivity, with slower alpha oscillations correlating with higher pain sensitivity [13].

Recent advances in neuromodulation techniques, particularly transcranial alternating current stimulation (tACS), have provided new opportunities for understanding and manipulating these oscillatory activities [2,4,18,24,32,42]. While tACS applied at theta, alpha, and beta frequencies—with or without additional modalities such as expectation modulation [2] or physical therapy [4], or as a standalone intervention [18,32,42]—has shown promising results in reducing pain, there remains a lack of sufficient understanding of the underlying mechanisms. The limited and contradictory reports on the efficacy of tACS for pain may be attributed to high interindividual variability and the widespread reliance on ‘one-size-fits-all’ tACS protocols with fixed frequencies, highlighting the need for individualized interventions.

Ahn et al. (2019) showed that 10 Hz tACS can enhance alpha oscillations in the somatosensory cortex of patients with chronic low back pain, correlating with pain relief. May et al. (2021) investigated the effects of tACS on tonic experimental pain in healthy participants, applying alpha and gamma tACS over somatosensory and prefrontal cortices. While their finding didn’t support that tACS can modulate tonic experimental pain, they suggested that optimizing stimulation parameters, including individualized stimulation frequencies aligned with each participant’s peak neural oscillations, could enhance efficacy. tACS at individual alpha frequency (IAF) has been shown to effectively entrain endogenous alpha oscillations [1,16,41], which might influence pain perception and associated neural dynamics [17,19,24,31,40].

We employed individualized tACS at IAF over the primary motor cortex (M1) of healthy participants to investigate its effects on pain perception and neural oscillations. M1 was chosen due to its central role in modulating corticospinal circuits and descending pain control pathways, as well as its frequent use as a brain stimulation target for chronic pain treatment according to a recent literature review [21]. We further used a discriminative approach to classify sensorimotor-related independent components (ICs), which were then utilized to reconstruct EEG signals and identify the SM-IAF. Sustained periodic thermonociceptive stimuli were delivered both before and after tACS. Behavioral responses—including stimulus intensity, painfulness rate, warm detection threshold (WDT), and heat pain threshold (HPT)—were recorded alongside EEG signals during thermal stimulation. Based on the proposed design using sustained periodic pain stimuli, our hypotheses were that tACS at the selected IAF: i) modulates pain perception by modulating alpha activity, and ii) alters phase-locked and pain-related ongoing oscillatory responses in the theta, alpha, and beta bands.

## 2. Method

### 2.1. Participants

Sample size: An *a priori* sample size calculation using G*Power [10], based on repeated measures ANOVA with a medium effect size (0.25), power of 0.95, and alpha level of 0.05, indicated a required sample of 36 participants.

We also conducted a simulation-based power analysis to determine the required sample size based on Linear Mixed Models (LMM) analysis with the regression formula: Measurement ∼ Condition * Time + (1∣Participant). Using pilot data of HPT, we assumed a baseline mean HPT of 46.7°C and a standard deviation (SD) of 4.3°C, applied for each Condition and Time point. A medium fixed effect size (*f*^2^=0.25) was used to model the condition × time interaction based on our hypothesis, corresponding to a mean difference of 2.25°C in the post active condition 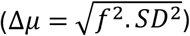. Simulations iteratively tested varying sample sizes to determine the smallest sample size required to achieve power of 0.9, at a significance level of 0.05. This approach resulted in a recommended sample size of 38 participants.

We recruited 40 participants to account for possible drop-outs; due to technical issues, data collection was incomplete for two individuals, resulting in a final dataset of 38 participants (19 females, 19 males; age: 28 ± 8.5). All experimental procedures were approved by the local Research Ethics Committee (AAHRPP-DSQ-037) and were performed in compliance with the Code of Ethics of the World Medical Association (Declaration of Helsinki). Prior to the experiment, all participants were informed about the procedure and potential side-effects and provided written informed consent. Participants received a financial compensation of 10 euros per hour.

The inclusion criterion for participants was being over 18 years. The exclusion criteria included pregnancy, a history of seizures or epilepsy, severe head injury, neurological or psychiatric illness, serious cardiac or respiratory conditions, metal implants in the head or scalp, cochlear implants, cardiac pacemakers, implanted neurostimulators (e.g., cortical, cerebral, vagus nerve, spinal cord), and implanted drug delivery devices.

### 2.2 Experiment design

We designed a within-subject double-blind experiment in which each participant completed two sessions (sham and active tACS), separated by approximately seven days, to minimize carryover effects (Fig. 1). The experiment followed a 2 × 2 design, with factors being Condition (sham, active) and Time (pre, post). The order of the sessions was counterbalanced across participants. Each session followed a structured protocol. The first step allowed participants to familiarize with thermonociceptive stimuli. This familiarization phase included three thermonociceptive stimuli with peak amplitudes similar to, but not identical to, those used in the main experiment. This approach aimed to help participants calibrate their pain ratings and reduce potential biases toward specific stimuli.

**Figure 1.**
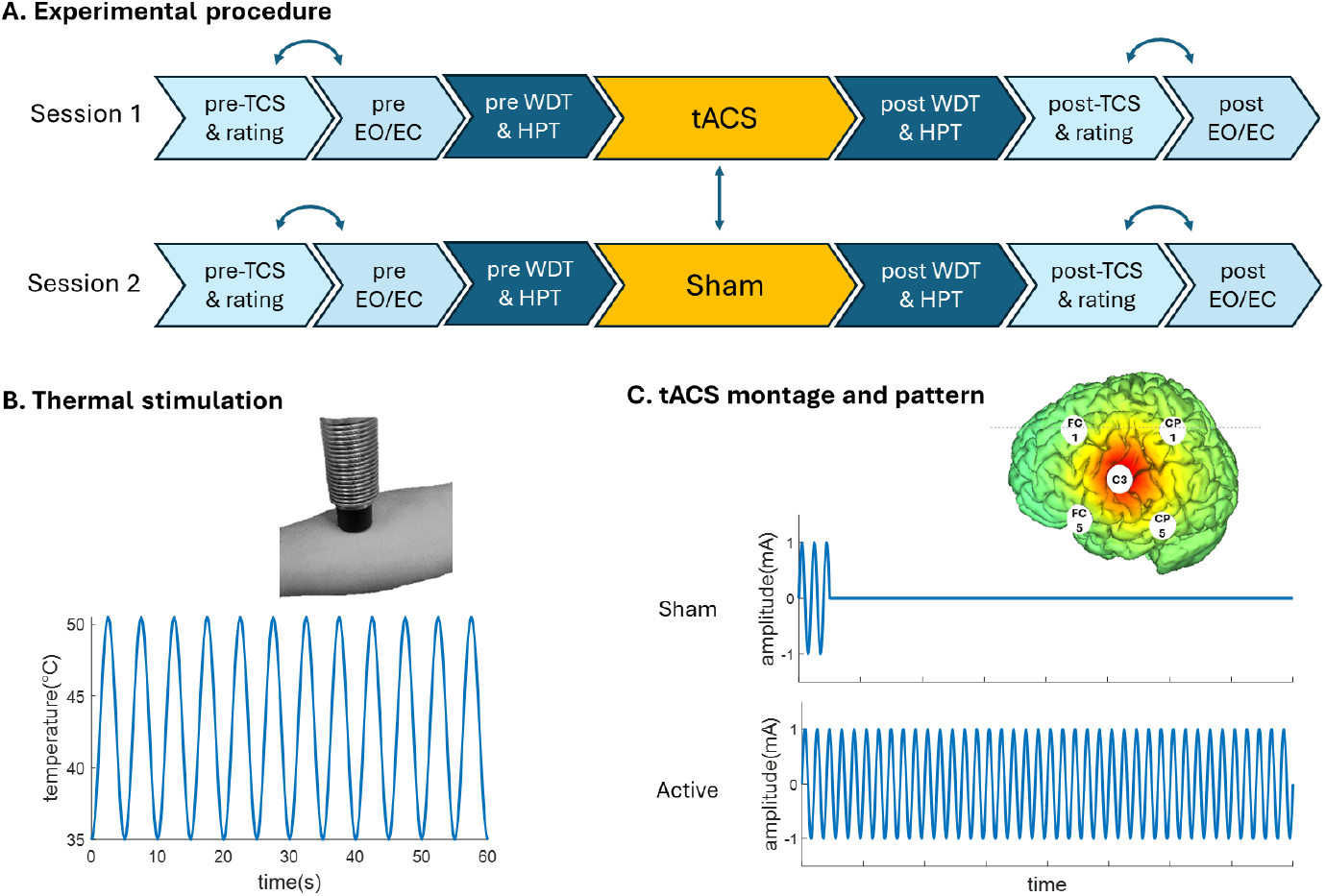
A. Timeline of the experimental protocol for active and sham tACS sessions. B. Illustration of thermal cutaneous stimulation applied to the forearm with a sinusoidal temperature waveform. C. High-definition 4×1 tACS montage targeting the primary motor cortex (M1), and stimulation patterns for sham and active conditions. Abbreviations: EO = eyes-open, EC = eyes-closed, TCS = thermal cutaneous stimulation.

Next, participants completed 4 minutes of rest during which EEG was recorded (2 minutes during eyes closed and 2 minutes during eyes open), and 10 trials of 1-minute sustained thermonociceptive stimulation. The order of these tasks was randomized and counterbalanced across participants, but remained consistent for pre- and post-tACS measurements. Furthermore, the same order was maintained for sham and active tACS sessions for each participant. The IAF was then computed using a custom MATLAB program (detailed in the IAF calculation section below), which took approximately 5 minutes. During this time, we also measured the warm detection threshold (WDT) and heat pain threshold (HPT). Participants then underwent 20 minutes of either sham or active tACS. Following tACS, the same procedures as before were repeated, beginning with WDT and HPT measurements. This was followed by 4 minutes of resting EEG (2 minutes eyes-closed, 2 minutes eyes-open) and 10 trials of 1-minute sustained periodic thermonociceptive stimulation.

### 2.3 Sustained thermonociceptive stimulation

To better understand how pain is related to neural oscillations, we used a long-lasting 1-minute, periodically-modulated stimulus (12 sin cycle with a frequency of 0.2 Hz) also used in previous studies^8^. In previous scalp EEG studies it has been shown that this type of stimulation can modulate ongoing neural oscillations at the frequency of stimulation [][][]. Moreover, the perception of sustained pain has been suggested to be associated with changes in the magnitude of modulated ongoing oscillations [][][]. Thermonociceptive stimuli were delivered on the volar forearm of the dominant arm of the participant using a thermal cutaneous stimulator (TCS, QST Lab), set with 15 micro Peltier elements (each with an approximate area of 7.7 mm^2^) whose temperature vary at rates of up to 300°C/s. The temperature of stimulation varied between a baseline temperature (35°C) and 50.5°C in a sinusoidal fashion. These parameters are known to elicit pain in the great majority of individuals^5,6,8^ and were also tested in pilot experiments before starting the main experiment. The stimulation probe was slightly displaced after each trial to limit sensitization.

### 2.4 Behavioral Measurements

After each of the 10 sustained periodic thermonociceptive stimulations, participants were prompted to rate their perception in response to a question displayed on the screen: “How intense did you perceive the stimulation?” (scale: 0–10 with 10 being the highest intensity imaginable). In addition, participants were required to respond to “Did you perceive the stimulation as painful?” (yes/no). The intensity ratings were averaged across the 10 trials, and the pain perception responses (0 for “no” and 1 for “yes”) were summed to provide a painfulness rate for each session.

For threshold measurements, participants were seated with their arm resting on a desk while a thermode (TCS, QST Lab) was applied to the forearm of their dominant hand. The temperature started at 27°C and increased at a rate of 1°C per second, with a maximum limit of 50°C. Without being able to see the temperature, participants were instructed to focus on their perception and press a button with their other hand as soon as they felt the slightest warming sensation (for warm detection threshold, WDT) and, subsequently, the first sensation of pricking or burning (for heat pain threshold, HPT). It was emphasized that the goal was to detect the initial sensation of heat pain, not to measure heat pain tolerance. WDT was measured before HPT to progress from less to more intense stimuli, reducing confounds such as sensitization or desensitization. Each measurement was repeated three times, and the average value was used.

### 2.6 EEG recording

EEG data were recorded using the same Starstim 32 system (Neuroelectrics, Barcelona, Spain) employed for tACS. For EEG recording, we used 32 Ag/AgCl electrodes based on the International 10-10 system with a DRL (Driven Right Leg) and CMS (Common Mode Sense) attached to the earlobe. EEG was recorded at a 500 Hz sampling rate during thermonociceptive stimulation and rest periods with eyes open and eyes closed, both before and after tACS. Electrode impedances were maintained below 20 kΩ. The recorded data were stored for offline processing.

#### 2.7.3 IAF determination (online processing)

The recorded signals were first preprocessed, including bandpass filtering (0.05–45 Hz) of signals recorded during TCS, EC, and EO using a 4th-order Butterworth filter and segmented into 1-minute trials. After detrending and mean removal, signals were re-referenced to the average of all channels. Wavelet ICA was then applied to automatically remove or suppress movement-related artifacts [6,20]. For left-handed participants (n=2), channels were symmetrically reordered across hemispheres (e.g., C4 for C3) to ensure consistent interpretation and analysis.

To measure the IAF, we employed an ICA-based method developed by Lebrun et al. [29] to dissociate vision-related alpha-band activity (VIS-alpha) and sensorimotor alpha-band activity (SM-alpha). An ICA was used to decompose the EEG signals recorded during the rest eyes-closed (EC), rest eyes-open (EO) and TCS-stimulation conditions into 31 components. The classification of ICs involved two main steps. First, using the average of the frequency spectra obtained in all three conditions (EC, EO, and TCS), potential alpha-band activity captured by each IC was identified by computing the peak frequency within the 7–13 Hz range using the center of gravity (CoG) formula:

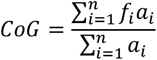

Where *f*_*i*_ is the i^th^ frequency bin, *a*_*i*_ is the spectral amplitude for *f*_*i*_, and n is the number of bins in the 7–13 Hz range. A 6 Hz frequency window centered on the CoG was then defined. Within this window, the mean amplitude and standard error of the mean (SEM: standard deviation of amplitudes divided by the square root of the number of bins) were computed separately for each condition. Confidence intervals (CIs) were derived as mean ± 1.97*SEM for each condition.

Based on the comparison of confidence intervals (CIs) across conditions, ICs were categorized as follows:

1. **Sensorimotor-IC (SM-IC)**: Components displaying a decrease in alpha amplitude during thermal cutaneous stimulation (TCS) without modulation by eyes-open (EO) or eyes-closed (EC) conditions.
2. **Visual-IC (VIS-IC)**: Components with higher alpha amplitude in EC compared to EO and TCS, but not sensitive to TCS.
3. **MIX-IC**: Components showing an increase in alpha amplitude in EC relative to EO and a decrease during TCS compared to EO.
4. **None**: Cases where the algorithm could not classify ICs into the above categories.

Figure 2 shows examples of SM-ICs and VIS-ICs under different recording conditions. EEG signals were reconstructed using ICs from each category, and the IAF was determined based on the center of gravity (CoG) within the 7–13 Hz range. The PAF at C3 (or C4 for left-handed participants) was designated as the IAF. Priority was given to PAF extracted from EEG reconstructed by SM-ICs, followed by MIX-ICs, then VIS-ICs. If no components matched these categories, a default frequency of 10 Hz was chosen for tACS stimulation. This approach is similar to the approach described by Gundlach et al. [15], where sensorimotor-related alpha frequencies were derived from recordings during passive somatosensory stimulation.

**Figure 2.**
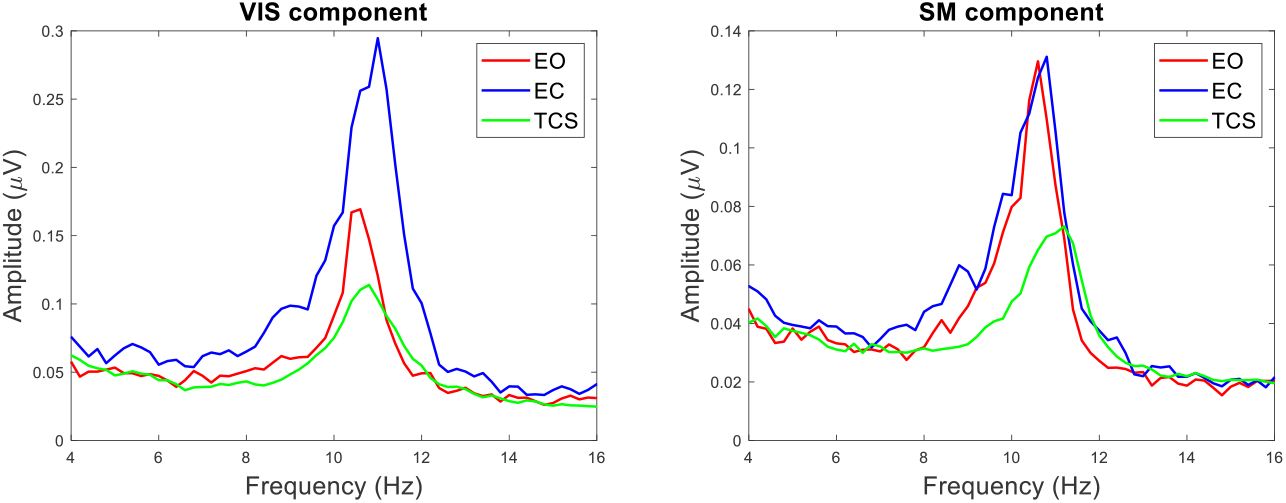
Classification ICs into VIS and SM categories based on their alpha-band activity. Frequency spectra showing the VIS-IC (left) and SM-IC (right) under different conditions: eyes-open (EO), eyes-closed (EC), and thermal cutaneous stimulation (TCS). The VIS component exhibits higher alpha amplitude during the EC condition compared to EO and TCS, whereas the SM component shows reduced alpha amplitude during TCS compared to EO or EC conditions.

### 2.5 Transcranial Alternating Current Stimulation (tACS)

tACS at the IAF was delivered for 20 minutes to the primary motor cortex (M1) using a 32-channel tES-EEG device (Starstim 32, Neuroelectrics, Barcelona, Spain). This device enabled EEG recording and electrical stimulation within a single setup. Once the neuroprene headcap was in place, EEG signals could be recorded or brain stimulation could be delivered to specific targets by selecting the corresponding electrodes. We applied a high-definition 4×1 montage to achieve focal stimulation (Fig. 1C) A sine-wave electrical current with a peak amplitude of 1 mA was delivered to M1 contralateral to the nociceptive stimulation. The center of stimulation was electrode C3 (or C4 for left-handed participants), surrounded by four electrodes positioned in a ring approximately 5 cm away at FC1, FC5, CP1, and CP5 (or FC2, FC6, CP2, and CP6 for left-handed participants) to enhance focality. The surrounding electrodes received 25% of the total current. We used Ag/AgCl based electrodes, each with a surface area of 3.14 cm^2^.

For sham stimulation, the protocol included a 30-second ramp-up followed by a 30-second ramp-down at the beginning of the session. At the end of each session, participants answered three questions. They rated the intensity of tingling or pricking sensations perceived on the scalp (0–10, with 10 being the most intense tolerable), and whether they believed they received continuous stimulation (0 = no, 1 = yes). The order of sham and active stimulation was randomized for each participant and uploaded to the stimulation device via MATLAB scripts. Both the participant and the operator were blinded to the stimulation protocol.

### 2.7 EEG data Analysis (offline processing)

#### 2.7.1 IAF offline analysis

In addition to the online calculation of IAF used to determine the frequency of stimulation, post-hoc analyses were conducted to compare IAF pre- and post-stimulation. The data were preprocessed as described in the online IAF calculation section, including bandpass filtering, re-referencing, and artifact removal. The same component classification procedure (SM-IC, VIS-IC, MIX-IC) was applied, and IAF was determined for each condition (pre- and post-stimulation) using the center of gravity (CoG) method. This allowed for a direct comparison of changes in IAF induced by tACS. To provide a clearer understanding of the processing workflow, a schematic representation of the preprocessing and analysis steps is presented in Figure 3.

**Figure 3.**
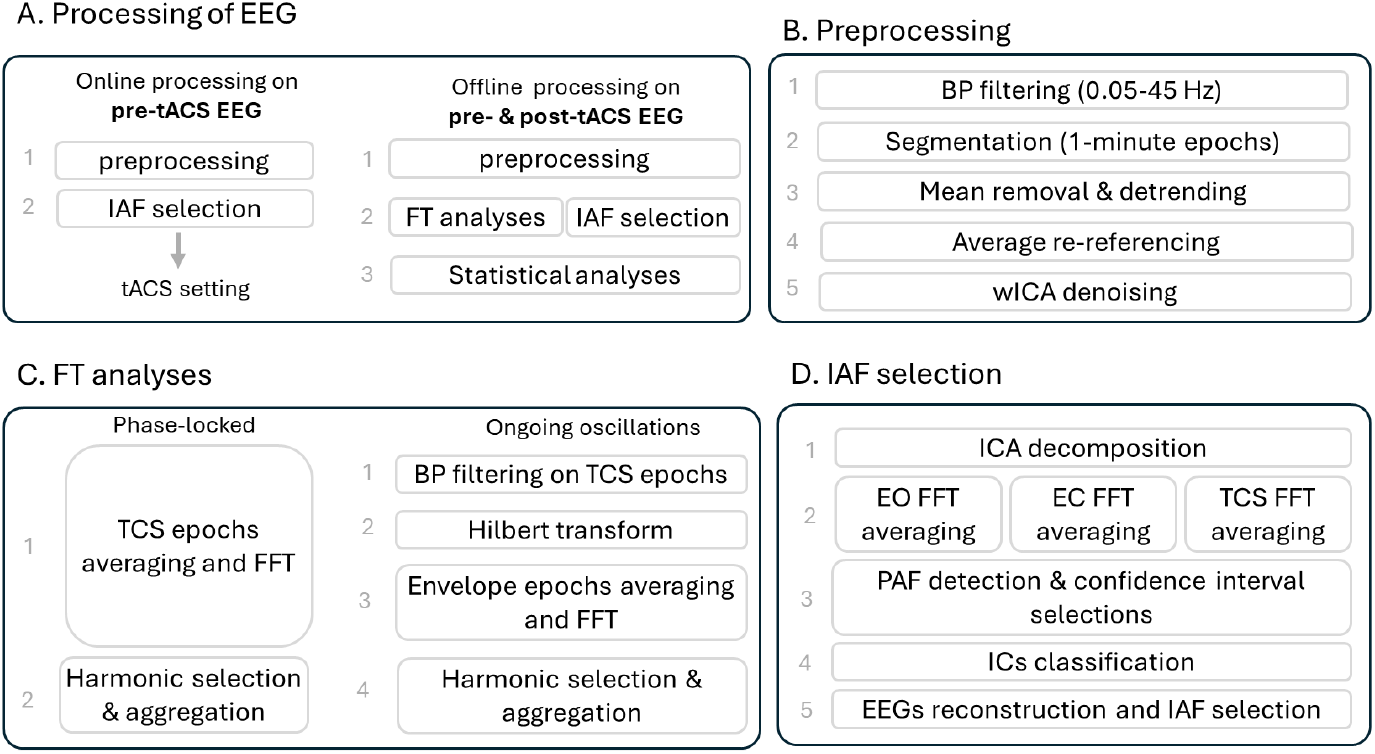
Overview of the EEG processing pipeline. A. Processing of EEG signals includes both online and offline analyses. Online processing of pre-tACS EEG involves preprocessing and IAF selection, which guides tACS settings. Offline processing of pre- and post-tACS EEG includes preprocessing, FT analyses, IAF selection, and statistical analyses. B. Preprocessing steps include bandpass filtering (0.05–45 Hz), segmentation into 1-minute epochs, mean removal, detrending, average re-referencing, and Wavelet ICA (wICA) denoising to remove artifacts. C. FT analyses are divided into two components: phase-locked responses (averaging TCS epochs, Fourier Transform, and harmonic selection/aggregation) and ongoing oscillatory activity (bandpass filtering, Hilbert transform for envelope extraction, averaging, and harmonic aggregation). D. IAF selection workflow, consisting of ICA decomposition, averaging FFT for EO, EC, and TCS conditions, detection of peak alpha frequency and confidence intervals, IC classification (SM-IC, VIS-IC, MIX-IC), and reconstruction of EEG signals to finalize IAF for tACS stimulation.

#### 2.7.2 FT analyses

We used periodic thermonociceptive stimuli to modulate neural ongoing oscillations at the frequency of stimulation, enabling us to analyze their modulation in relation to pain perception. To analyze these oscillatory responses, we used frequency-tagged analysis, a method that isolates neural activity synchronized with the frequency of the periodic stimuli allowing us to track amplitude-specific changes in the brain’s response to thermonociceptive input.

We analyzed both the phase-locked response at the frequency of stimulation, and the modulation of ongoing oscillatory activity in the different frequency bands. The data were initially preprocessed as described in the above section on IAF calculations. For the phased-lock responses, after preprocessing and segmentation of EEG signals into the 1-minute epochs, we averaged data over epochs for each participant and each condition and then transformed them to the frequency domain using a Fourier transform. Using this frequency-domain analysis approach we expect to see distinct peaks in the EEG frequency spectrum at the stimulation frequency and its harmonics. To better estimate the response amplitude at each frequency bin, we subtracted the average amplitude of the signal measured at neighboring frequencies (−0.026 to -0.065 Hz and 0.026 to 0.065 Hz) to reduce residual noise [27].

The signal-to-noise ratio (SNR) for each harmonic was defined as the ratio of harmonic power to baseline noise power and calculated as:

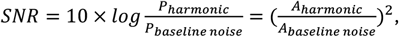

where *A*_*harmonic*_ represents the amplitude at each harmonic, and *A*_*baseline noise*_ is the average amplitude of the signal at neighboring frequencies (−0.026 to -0.065 Hz and 0.026 to 0.065 Hz), as defined above. We aggregated harmonics whose SNR, averaged across all channels, was greater than zero. Aggregated amplitudes were used for statistical tests.

For analyzing the ongoing oscillatory response to periodic stimuli, a similar procedure was applied. After preprocessing and segmenting the EEG signals into epochs, the signals were bandpass filtered into theta (4-8 Hz), alpha (8-13 Hz), and beta (13-30 Hz) frequency bands consistent with COBIDAS guidelines [38] using Butterworth bandpass filters. The signal envelopes within each band were then extracted using the Hilbert transform. Finally, the envelopes were averaged across epochs for each participant and condition and transformed into the frequency domain using the Fourier transform. Residual noise correction and SNR calculation for harmonics were performed similarly to the approach used previously for phase-locked responses.

### 2.8 Statistical analyses

Statistical analyses were conducted using LMMs with the fitlme function in MATLAB (2023a, The MathWorks). The model was fitted using restricted maximum likelihood (REML), ensuring robust estimation of both fixed and random effects.

Residuals of the LMM were checked for normality using both a Q-Q plot and the Shapiro-Wilk test. If residuals were not normal, skewness was evaluated. For skewness values greater than 0.5, alog transformation was applied to normalize the data. Outliers were then identified using Cook’s distance, and the sample with the highest Cook’s distance was removed if it exceeded a threshold of 3 standard deviations of the distance. The model was then refitted, and residuals were reassessed. If residuals still showed non-normality, the outlier removal process was repeated. This procedure was limited to a maximum of 5% of the dataset to ensure no more than 5% of data was removed. Normality was achieved in all cases before exceeding the 5% data removal limit.

#### 2.8.1 Behavioral response

To examine the effect of tACS on behavioral response we performed four LMM analyses with the following model: Measurement ∼ Condition * Time + (1 ∣ Participant), where Measurement refers to the dependent variable, which varied based on the analysis. It could represent, HPT, WDT, painfulness rate, or intensity ratings. Condition (sham vs. active) and Time (pre-vs. post-stimulation) were included as fixed effects, along with their interaction. Random intercepts were specified for Participant to account for between-subject variability.

#### 2.8.2 Phase-locked and ongoing oscillations responses

As described earlier, aggregated amplitudes of harmonics with SNR higher than zero were used for statistical tests. We used a permutation-based significance test to identify the channel with the most significant modulations across all conditions, testing whether the peak aggregated amplitudes at each channel were significantly greater than zero [23]. First, a one-tailed Wilcoxon signed-rank test was applied to each channel to assess whether its values differed significantly from zero. To account for multiple comparisons across the four conditions (sham vs. active, pre-vs. post-stimulation), we applied a Bonferroni-corrected threshold of α=0.05/4 = 0.0125. A null distribution was generated by randomly flipping the signs of the data in 2000 permutations. Channels were regarded as significant if their p-values were below the threshold and their count exceeded the 95th percentile of the null distribution, ensuring robust control of false positives. We selected the most significant channel with the highest test statistic for further analyses.

To see the tACS effect on phase-locked responses, we performed LMM analyses with the following formula: Measurement ∼ Condition * Time + (1 ∣ Participant), where Measurement refers to the aggregated amplitude of the phase-locked FT response at the most significant channel. For ongoing oscillations, we used the same model (Measurement ∼ Condition * Time + (1 ∣ Participant)) and analyses, but here Measurement referred to the aggregated amplitude of the ongoing oscillations’ frequency-tagging (FT) response in the theta, alpha, or beta frequency bands, with the most significant channel determined separately for each band.

#### 2.8.3 IAF data

To evaluate the IAF data, we performed LMM analyses using the same model, with IAF as the dependent variable and Condition, Time, and their interaction as fixed independent factors, and Participants as a random factor.

## 3. Results

### 3.1. Behavioral responses

The results of behavioral measurements for all participants at different conditions and time points are shown in Figure 4. The LMM results for behavioral measures indicate a significant decrease in WDTs (F=9.5; p = 0.002) and HPTs (F=11.7; p= 0.0007) for both post-active and post-sham conditions, with no significant interaction between time and condition (F=0.8; p=0.35 for WDT; F=0.1; p=0.78 for HPT). Additionally, participants showed a significant increase in perceived stimulus intensity during sustained periodic stimulation (F=22.7; p<0.0001), as well as an increase of painfulness rate (F=17.9; p<0.0001). These consistent decreases in thresholds and increases in perceived intensity ratings may reflect mild sensitization resulting from sustained thermal stimulation.

**Figure 4.**
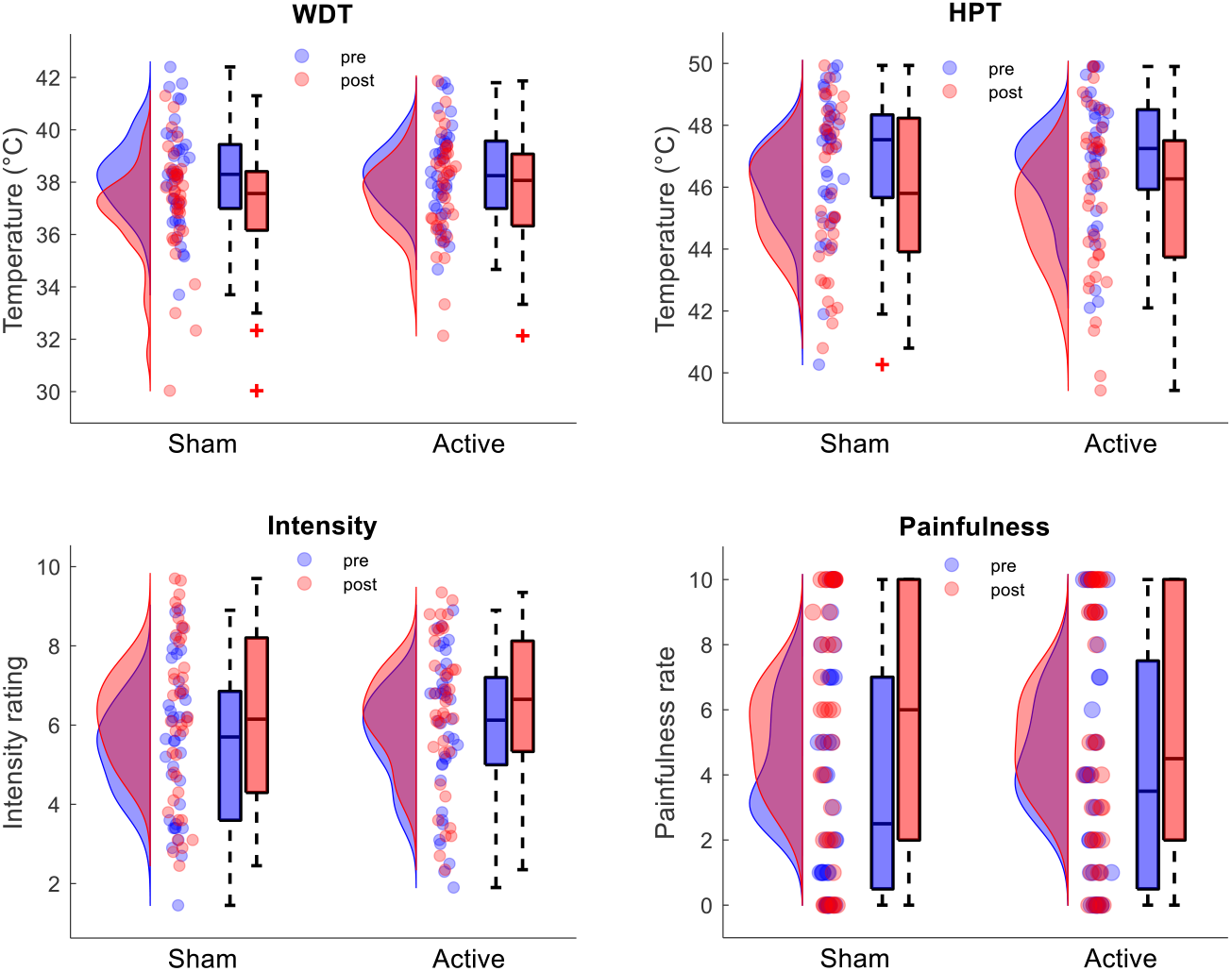
Variations in WDT, HPT, stimulus intensity ratings, and painfulness rates across different conditions and time points for all participants. LMM results for all measurements show a significant effect of time for all measurements.

Figures also show that the average decrease in WDT and HPT, as well as the increase in intensity and painfulness rate, were less pronounced in the active tACS condition compared to sham. The average decrease in WDT and HPT was 1.1°C for sham and 0.6°C for tACS, while the average increase in painfulness rate on a 0-10 scale was 0.16 for sham and 0.11 for tACS. For intensity on a 0-10 scale, the average increase was 0.61 for sham and 0.51 for tACS. However, statistical analyses indicate that these differences did not result in significant interactions (see Table S1).

### 3.2 Neural responses to sustained periodic thermonociceptive stimulation

The results of the Wilcoxon signed-rank test showed that the strongest phase-locked periodic modulation across all conditions and time-points occurred at FC2. For ongoing oscillations, the test revealed that the strongest periodic modulation was at Cz for the theta frequency band and at C3 for the alpha and beta frequency bands, respectively (Fig. 5). The results of each condition and time points can be found in Figure S2.

**Figure 5.**
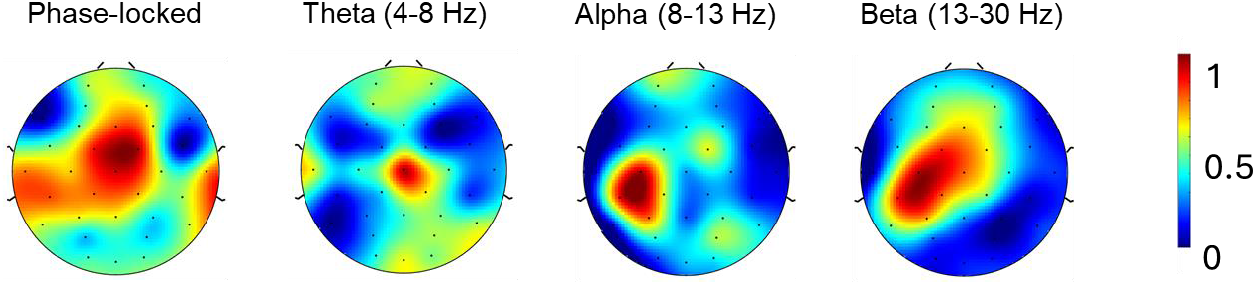
Topographical maps showing the strongest phase-locked activity and ongoing oscillatory modulations in response to periodic thermonociceptive stimulation considering all conditions and time points together. The phase-locked response is localized at FC2, while ongoing oscillatory activity shows the strongest modulations at Cz for the theta band (4–8 Hz) and at C3 for the alpha (8–13 Hz) and beta (13–30 Hz) bands. Color scales represent Wilcoxon test statistics.

In Figure 6, the frequency spectra at the most significant electrodes, averaged across participants, illustrate the neural phase-locked and oscillatory responses to periodic thermonociceptive stimulation under different conditions and time points. The spectra show decreased amplitudes in post-stimulation compared to pre-stimulation, with a more pronounced reduction in the phase-locked response. LMM analyses for phase-locked responses showed that there was a significant time effect (F=5.0; p=0.03), but no condition effect or interactions between time and condition (see Table 1). The analyses of ongoing oscillations at theta, alpha, and beta showed no significant time, condition, or interaction effect on periodic modulations (Table 1). Figure 7 shows the distribution and boxplots of aggregated harmonic amplitudes for phase-locked responses, theta, alpha, and beta bands across sham and active conditions. A decrease in amplitudes post-stimulation compared to pre-stimulation is observed for phase-locked responses, while other frequency bands do not show a consistent change.

**Table 1.** Results of statistical tests for phase-locked and ongoing oscillations at the channel with the most significant amplitude modulations by thermonociceptive stimulation: phase-locked (FC2), theta (Cz), alpha (C3), and beta (C3).

|  | Phase-locked (FC2) | Theta (Cz) | Alpha (C3) | Beta (C3) |
| --- | --- | --- | --- | --- |
| Condition | F=0.5, p=0.47 | F=0.0, p=0.92 | F=1.9, p=0.17 | F=0.2, p=0.69 |
| Time | **F=5.0, p=0.03** | F=0.15, p=0.69 | F=0.1, p=0.80 | F=3.0, p=0.09 |
| Interaction | F=0.4, p=0.54 | F=0.1, p=0.76 | F=0.2, p=0.67 | F=2.0, p=0.16 |

**Figure 6.**
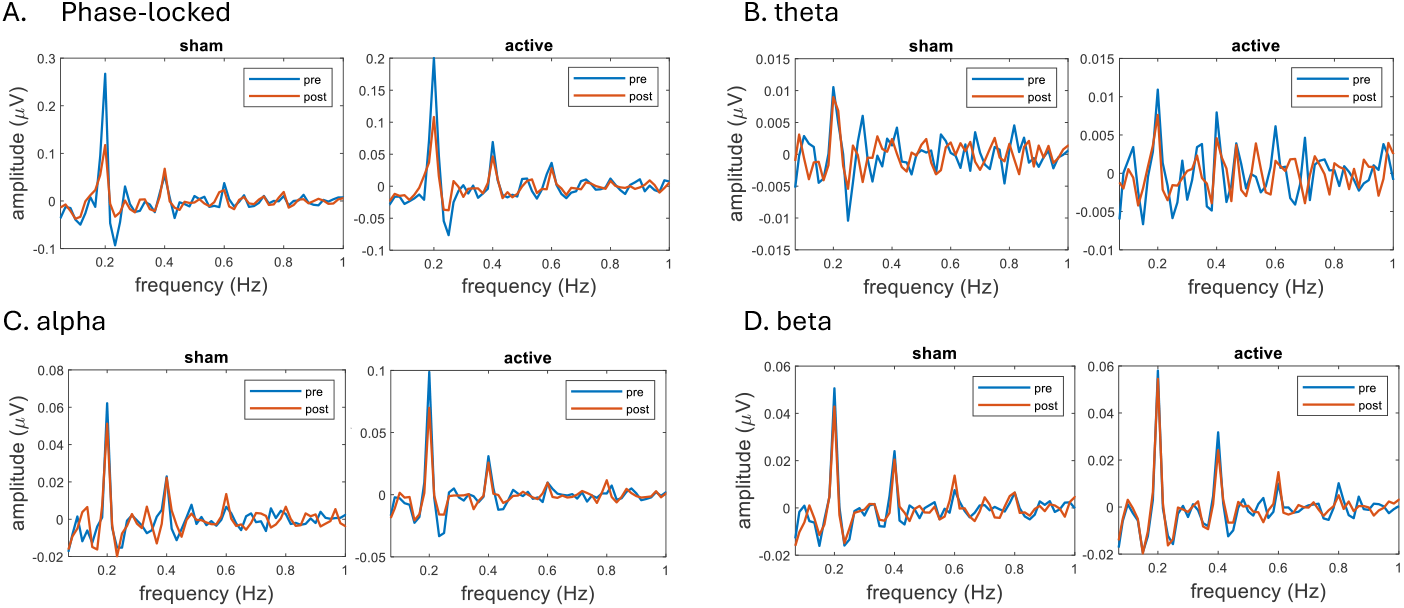
Frequency domain representation of phase-locked and oscillatory activity in response to periodic thermonociceptive stimulation averaged across all participants. (A) Phase-locked, (B) theta-band oscillations, (C) alpha-band oscillations, and (D) beta-band oscillations are shown separately for sham and active conditions. Pre-stimulation (blue) and post-stimulation (red) spectra highlight changes in amplitude across conditions.

**Figure 7.**
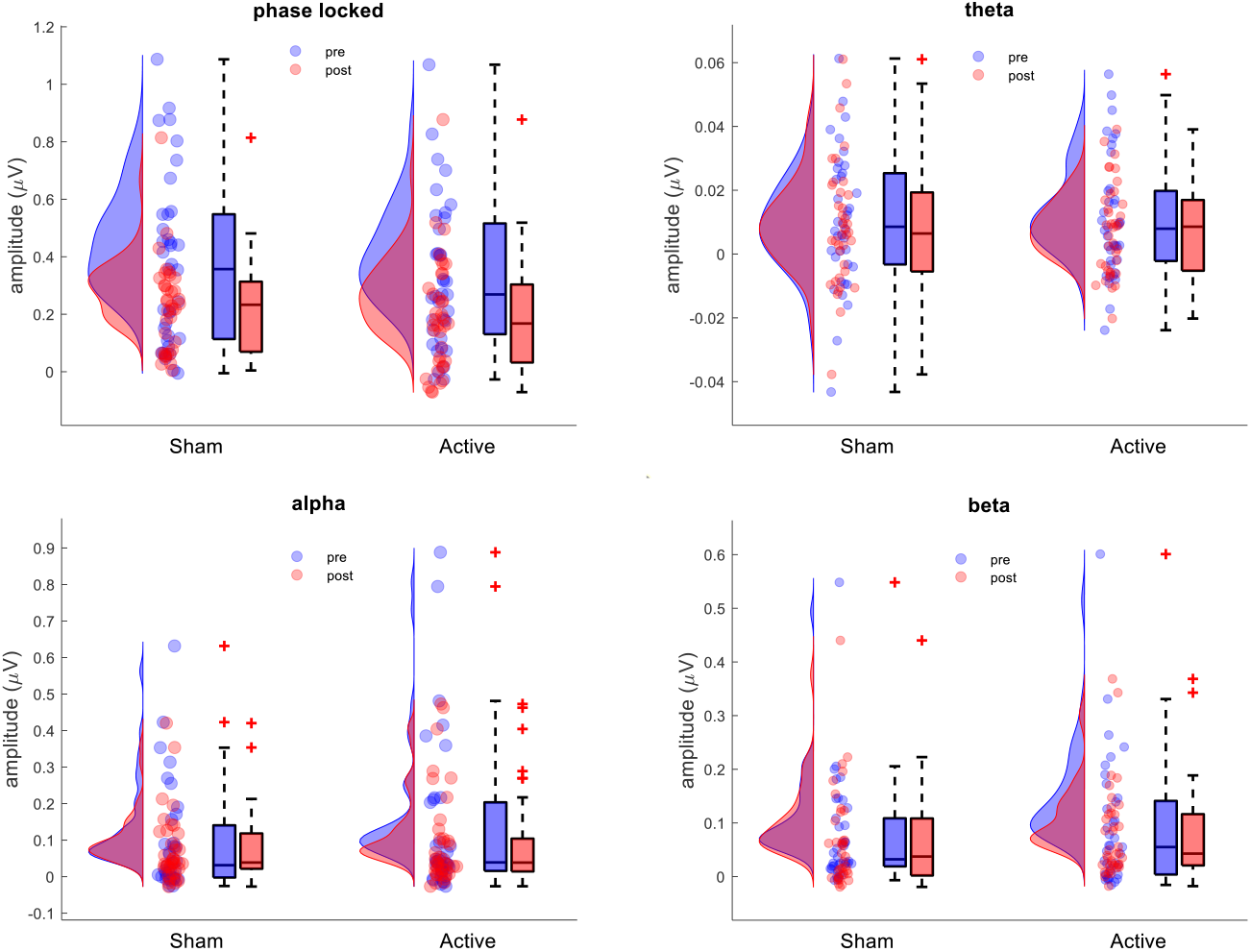
Distributions and boxplots of amplitudes across participants for phase-locked responses (top left), theta band (top right), alpha band (bottom left), and beta band (bottom right) during sham and active conditions.

To examine the relationship between variations in behavioral responses and phase-locked responses, we performed a correlation analysis. However, no significant correlation was found between the variation in phase-locked amplitudes and behavioral measurements (Fig. 8).

**Figure 8.**
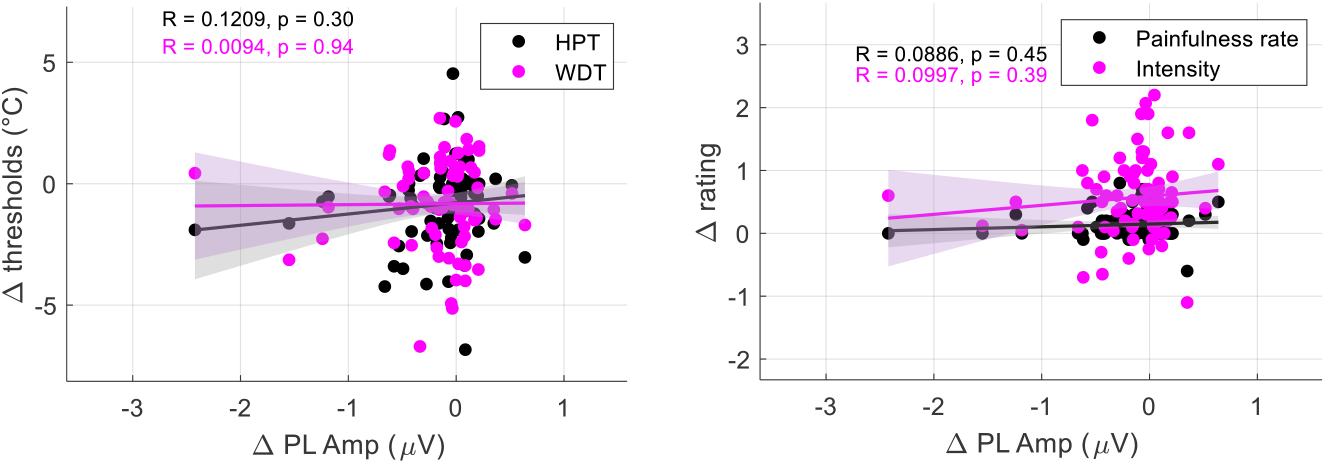
Correlation analyses between changes in phase-locked amplitudes (Δ PL Amp, µV) and changes in behavioral measures across time points, regardless of conditions (sham or active). Δ PL Amp = post phase-locked amplitudes - pre phase-locked amplitudes, Δ thresholds = thresholds post – thresholds pre, and Δ rating = post rating– pre rating.

### 3.3 IAF Analyses

In 26 subjects SM-IAF was detected, in 3 subjects VIS-IAF was detected and in 7 subjects MIX-IAF was detected; in the remaining 2 subjects, none of these three specifics ICA-based IAF were detected, so 10 Hz was selected as a default tACS frequency during the experiment. Figure 9 shows how different channels contributed to extract SM-IC, MIX-IC, and VIS-IC.

**Figure 9.**
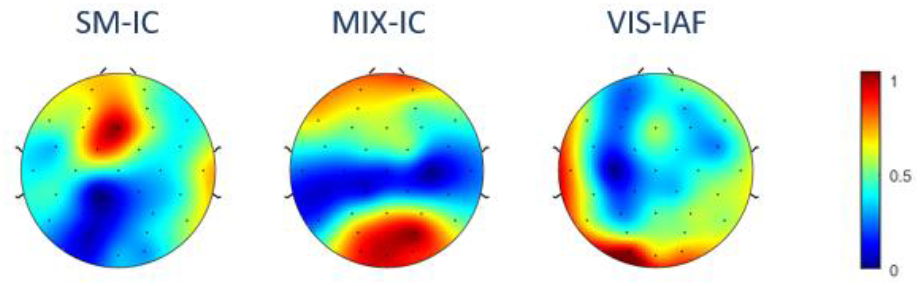
Spatial weight topography for selected IAF ICs, showing the contribution of each channel to the components mixing matrix derived from ICA.

In Figure 10, the average alpha spectral pattern across participants shows greater fluctuations post-sham compared to pre-sham, than the fluctuations observed post-active compared to pre-active. In particular, during SM-IAF tACS, the IAF remained more stable, with less deviation from its pre-stimulation position, whereas the IAF shifted to the left (lower frequencies) during sham for the same participants. The distribution of selected IAF across participants is presented in Figure 11. Comparing overall selected IAF at C3/C4 location across conditions and time using LMM showed no significant changes across conditions or time points. When considering only SM-tACS-IAF, a marginal time effect (F = 3.9, p = 0.05) and a trend toward an interaction for the post-active condition (F = 3.2, p = 0.07) were observed.

**Figure 10:**
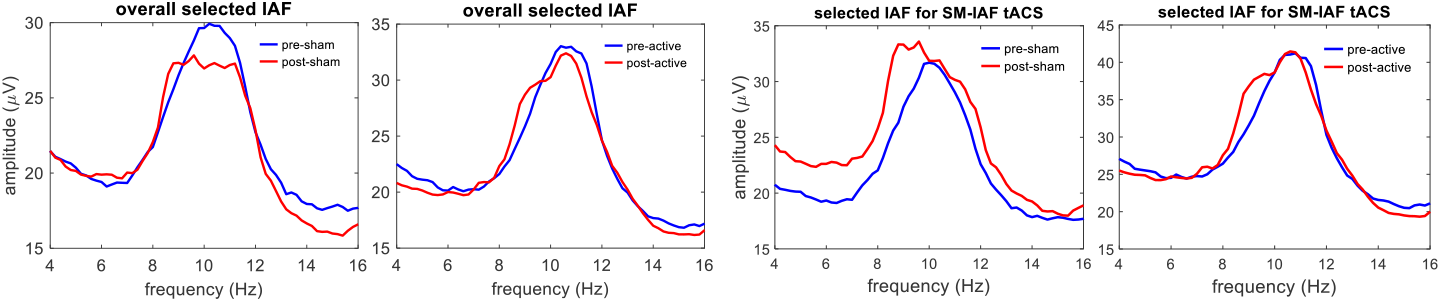
Alpha spectral pattern averaged across all participants for each condition. The most clear shift can be seen for post-sham SM-IAF compared to pre-sham SM-IAF.

**Figure 11:**
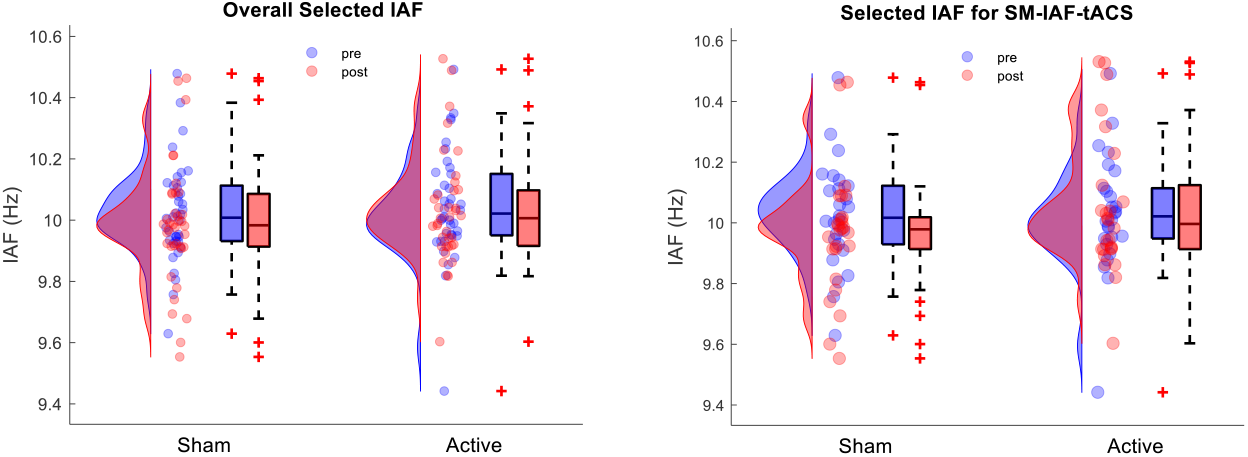
Selected IAF at the C3/C4 location across experimental conditions and time points. LMM analysis revealed no significant differences between conditions (p = 0.38), time points (p = 0.14), or their interaction (p = 0.51) for the overall selected IAF across all participants (n = 38). However, for those who received tACS at SM-IAF (n = 26), a marginal time effect (F = 3.9, p = 0.05) and a trend toward an interaction for the post-active condition (F = 3.2, p = 0.07) were observed.

To investigate the relationship between variations in pain perception and SM-IAF, we performed correlation analyses as shown in Figure 12. The results revealed that only HPT showed a marginally significant correlation with SM-IAF variations (R = 0.28, p = 0.05).

**Figure 12.**
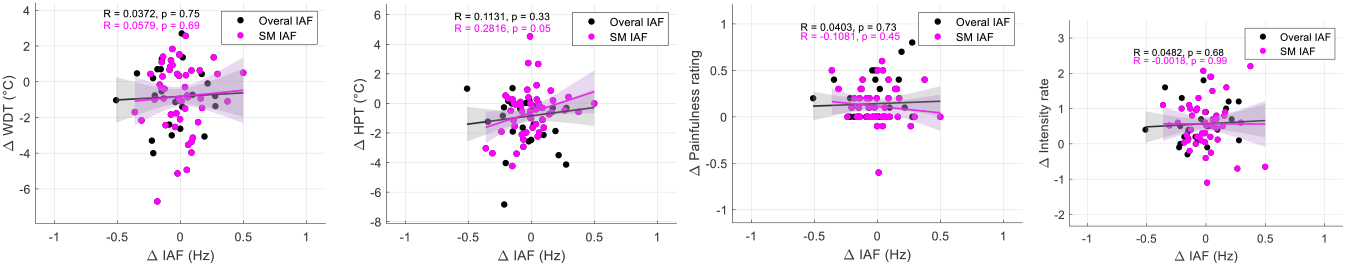
Correlation analyses between changes in pain perception measures (WDT, HPT, painfulness rating, and intensity rating) and changes in IAF (overall IAF and SM-IAF). Only HPT showed a marginally significant correlation with SM-IAF (R = 0.28, p = 0.05).

Figure 13 provides a detailed view of how tACS at SM-IAF affects HPT. Correlation analyses reveal that a significant correlation exists between SM-IAF variations and HPT for the sham condition (R = 0.44, p = 0.03), indicating that a decrease in IAF is associated with a decrease in HPT. However, no such correlation is observed for the tACS condition.

**Figure 13.**
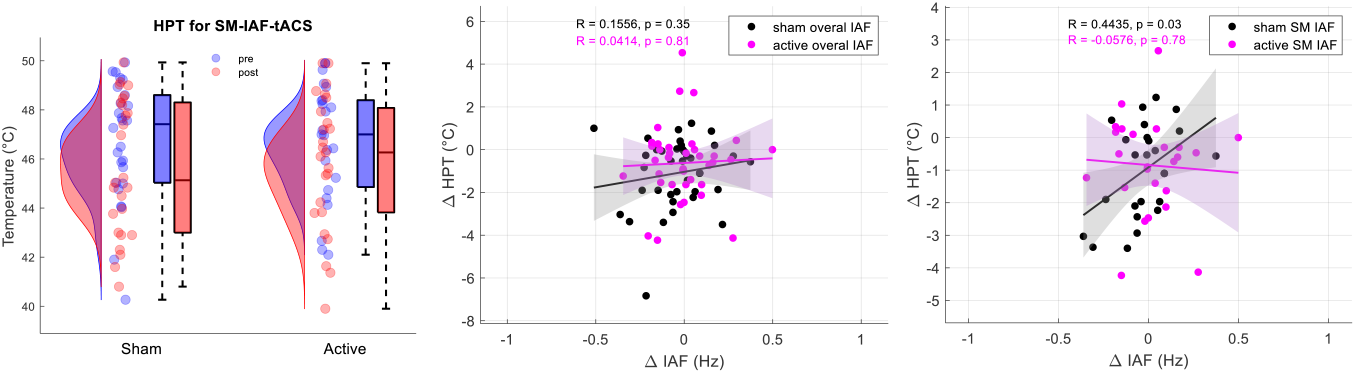
HPT relationship with IAF. Left: Distribution of HPT (°C) pre- and post-stimulation across all conditions for participants who received SM-IAF tACS. A significant time effect was observed (F = 9.3, p = 0.003), but no interaction (p = 0.47) or condition effect (p = 0.71). Middle and right: Correlation analyses between changes in IAF (Δ IAF) and changes in HPT (Δ HPT) for overall IAF (middle) and SM-IAF (right). A significant correlation between Δ SM-IAF and Δ HPT was found for the sham condition (R = 0.44, p = 0.03), indicating that a decrease in IAF is associated with a decrease in HPT, whereas no such correlation was observed for the tACS condition.

### 3.4. Exploring the effects of SM-IAF and sex on behavioral response to the tACS

We performed exploratory analyses to investigate how sex may influence the behavioral response to tACS, While the sample size for each subgroup (female/male/SM-IAF) is not sufficient to draw definitive conclusions, these analyses were intended to identify potential trends and define directions for future research. The results in Figure 14 (with statistical tests results in Tables S1 & S2) indicate that, in general, the time effect on behavioral measures was stronger in females than in males. Specifically, no significant time effect was observed for HPT in males, whereas a significant time effect was found for HPT in females. Additionally, a trend was observed in females, suggesting greater post-active tACS effects compared to post-sham conditions.

**Figure 14.**
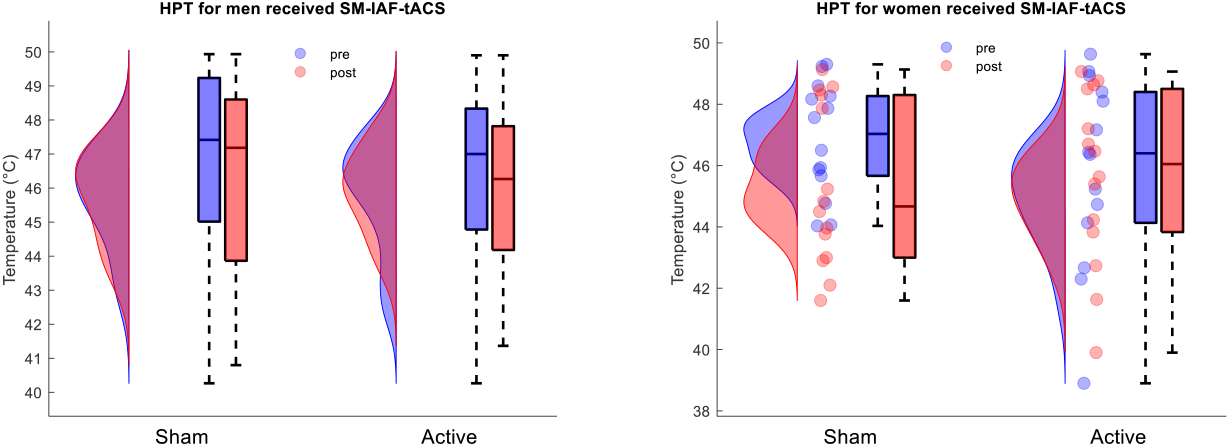
HPT distribution for men (left) and women (right) who received tACS at SM-IAF (n = 12 for men, n = 14 for women).

## 4. Discussion

In this study, we examined how individualized alpha-tACS applied to M1 affects neural oscillations and behavioral responses to sustained thermonociceptive stimuli in healthy participants. In this section, we discuss each of our hypotheses based on the observed results of previous section.

### 4.1 Individualized tACS and pain perception

Although the same thermal stimulation was delivered pre- and post-tACS, we observed an overall increase in pain perception and intensity ratings in both the sham and active conditions. The decreased WDT and HPT values in the post-tACS phase suggest mild sensitization of the forearm, which could also account for the increased ratings of sustained thermonociceptive stimuli.

While the average values showed that tACS resulted in a less pronounced decrease in WDT and HPT, as well as a smaller increase in intensity and painfulness ratings compared to sham, these differences were not statistically significant. Nevertheless, the consistency of these trends across the different behavioral measures suggests a potential modulatory effect of tACS on nociception processing and pain perception.

As a potential source of variability in pain perception, we conducted an exploratory analysis to examine the influence of sex on tACS modulation of behavioral responses. Overall, the comparison of post-phase to pre-phase revealed a more pronounced effect in women than in men, with HPT showing the most significant decrease over time in women and no significant change in men. Additionally, there was a trend for an interaction effect, indicating that females responded better to tACS at SM-IAF. Sex and gender differences in pain perception and sensitivity have been widely documented, with females generally showing higher pain sensitivity compared to males [37,44,45]. These differences may stem from hormonal, immune, and structural variations in nociceptive pathways. Regarding tACS or transcranial electrical stimulation (tES) in general, while research on sex differences is limited, anatomical differences, such as skull thickness and brain morphology, along with physiological factors like hormonal fluctuations, can influence the distribution and efficacy of electrical current during stimulation [5,22,51,52]. Adapting tES protocols to account for these factors could enhance the effectiveness of neuromodulation across diverse populations. These factors are also likely contributors to individual variability in IAF [3], further emphasizing the need for personalized approaches.

### 4.2 Neural phase-locked and ongoing oscillations responses to tACS

Frequency-tagging analysis revealed significant phase-locked modulations in frontal-central electrodes in response to periodic thermonociceptive stimuli, as well as in central and posterior-central areas for theta, and in central and parietal regions contralateral to the dominant arm for alpha and beta, consistent with previous findings in the literature [8,23]. Statistical tests on amplitude variations at the frequency of interest (0.2 Hz) and its harmonics showed no significant effects of time or condition on ongoing neural oscillations during sustained thermonociceptive stimulation. However, phase-locked responses significantly decreased in the post-stimulation phase compared to the pre-stimulation phase, regardless of condition. While no tACS effect was found on PL amplitude, the reduction in the “post” phase may be linked to habituation to the periodic aspect of the stimuli [26,36]. Moreover, the lack of observable tACS effects on oscillatory activities could be attributed to suboptimal stimulation parameters, as suggested by exploratory analyses (see Supplementary Table S2) showing marginal tACS effects when using SM-IAF. This highlights the need for more personalized and optimized stimulation protocols, along with sufficient sample sizes for each subgroup.

### 4.3 tACS effects on IAF and the relationship between IAF and pain perception

SM-IAF was identified using a discriminative ICA approach applied to data from different conditions. When SM-IAF could not be determined, other IAF types (MIX-IAF, VIS-IAF, or 10 Hz) were selected, which may influence the observed relationships. Comparing overall selected IAF values revealed no significant change in overall selected IAF; however, SM-IAF significantly decreased in the post phase compared to the pre phase regardless of condition. Correlation analysis showed a positive relationship between ΔHPT and ΔSM-IAF, indicating that higher sensitivity is associated with slower alpha frequencies. When analyzing the correlation between ΔSM-IAF and ΔHPT separately for the active and sham conditions, a significant correlation was observed only in the sham condition, with no correlation in the active condition.

We observed a positive correlation between ΔHPT and ΔSM-IAF, consistent with the relationship between IAF and pain perception observed in chronic pain patients [14,43,49,50] and some experimental pain studies [11–13]. In general, the relation between IAF and pain perception can be dependent on several factors including the type of pain (in this case, significant correlations were observed during threshold measurements but not sustained periodic thermonociceptive stimuli), the type of IAF (here, SM-IAF rather than overall selected IAF), and experimental conditions (e.g., tACS disrupted the significant correlation observed in sham experiments).

Valentini et al. highlight that while IAF can differentiate between painful and neutral sensations, it does not reliably index the affective unpleasantness of pain [47]. Their findings suggest caution in using IAF as a pain biomarker, emphasizing that methodological and statistical factors significantly influence alpha modulations and their interpretation. Additionally, May et al. assessed the predictive value of IAF for sensitivity to brief experimental pain, finding that variations in pain perception were not consistently predicted by IAF, indicating that the relationship between IAF and pain may not generalize to all types of pain [33]. These studies highlight the complexity of the relationship between IAF and pain perception, suggesting that while IAF holds promise as a biomarker for pain sensitivity, its applicability may vary across different pain modalities and individual differences. Ongoing research aims to deepen the understanding of how IAF variability impacts pain perception and its potential for guiding personalized neuromodulation strategies.

Alpha power analyses can be further refined and parameterized, for instance, by using the FOOOF algorithm [9], to separate periodic and aperiodic components, to define its bandwidth and power beyond just the peak frequency. This allows for more individualized strategies. In exploratory analyses (Fig. S3), we observed that the alpha spectrum curve during TCS was wider compared to eyes-open and the alpha spectrum had less amplitude compared to eyes-closed resting condition. Thes finding highlights differences in the alpha spectrum under various conditions, which could inform the development of more tailored stimulation strategies to optimize neuromodulatory outcomes, consistent with principles observed in Arnold tongue dynamics [16,41,46,48,53].

### 4.4 Concluding remarks and future directions

While the current study does not reveal a statistically significant effect of tACS on pain perception, the consistent trend of decreased pain perception and smaller PAF changes in the active condition compared to sham, alongside marginally significant effects observed in specific groups (e.g., SM-IAF tACS on women), suggests a potential reduction in sensitization following tACS. This highlights the potential for future studies to explore tACS effects on pain using individualized strategies that account for more individual variabilities and other condition-or state-dependent factors. While we used IAF as a personalization parameter, other tACS parameters, such as amplitude and montage, can also be personalized. Such strategies can be realized by building a mathematical model that incorporates several potential biomarkers and identifying model parameters through increased data collection to capture as many variabilities as possible.

## Acknowledgments

YF is supported by a MIS grant from the FNRS, and GL is supported by FNRS and Fondation Médicale Reine Elisabeth (FMRE) grants. We wish to thank all the members of the Nocions Lab (http://nocions.org) for the useful discussions.

## Author contributions

Conceptualization and Methodology: YF, GL; Data curation: YF, FD; Data Analyses: YF, GR; Writing - original draft: YF, GL; Writing – Review & Editing: YF, FD, GR, GL; Supervision, Project administration, and Finding acquisition: GL

## Conflicts of interest

The authors declare no competing financial interests or conflict of interest.

## Notes

### Competing Interest Statement

The authors have declared no competing interest.

