## Supplementary Material for "Individualized Alpha-tACS for Modulating Pain Perception and Neural Oscillations: A Sham-Controlled Study in Healthy Participants"

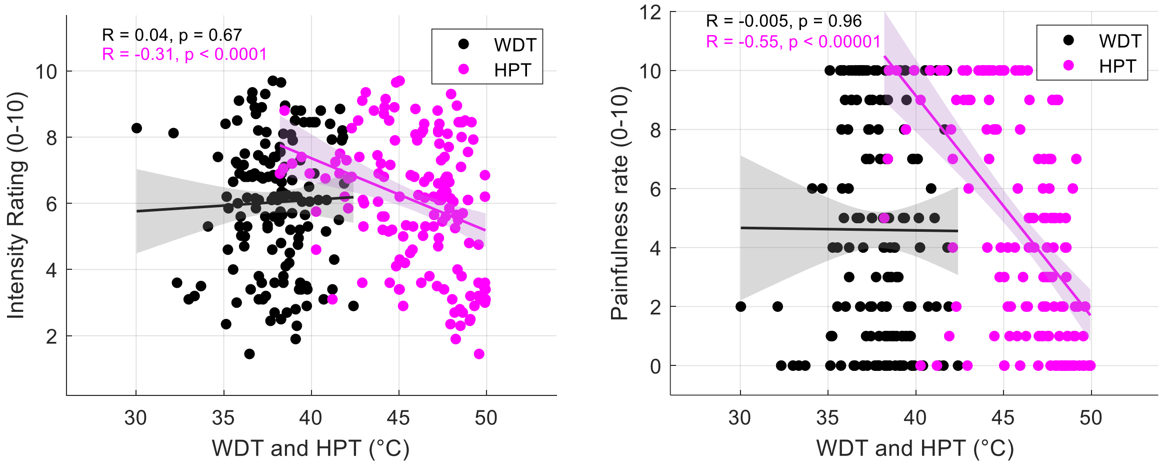


Figure S1. Correlation between thermal perception thresholds (WDT and HPT) and perceived intensity and painfulness rates. The left panel shows the relationship between WDT/HPT and intensity rating (0–10), while the right panel shows the relationship between WDT/HPT and painfulness rate (0–10). Linear regression lines and their respective confidence intervals are overlaid for each group. no significant correlations between WDT and intensity (R=0.04, p=0.67) or painfulness (R=-0.005, p=0.96) but significant correlations between HPT and intensity (R=-0.31, p<0.0001) and painfulness rate (R=-0.55, p<0.00001) were observed. The HPT high correlation with painfulness rate during sustained periodic thermonociceptive stimuli underscores the importance of using HPT as a primary measure in pain modulation research

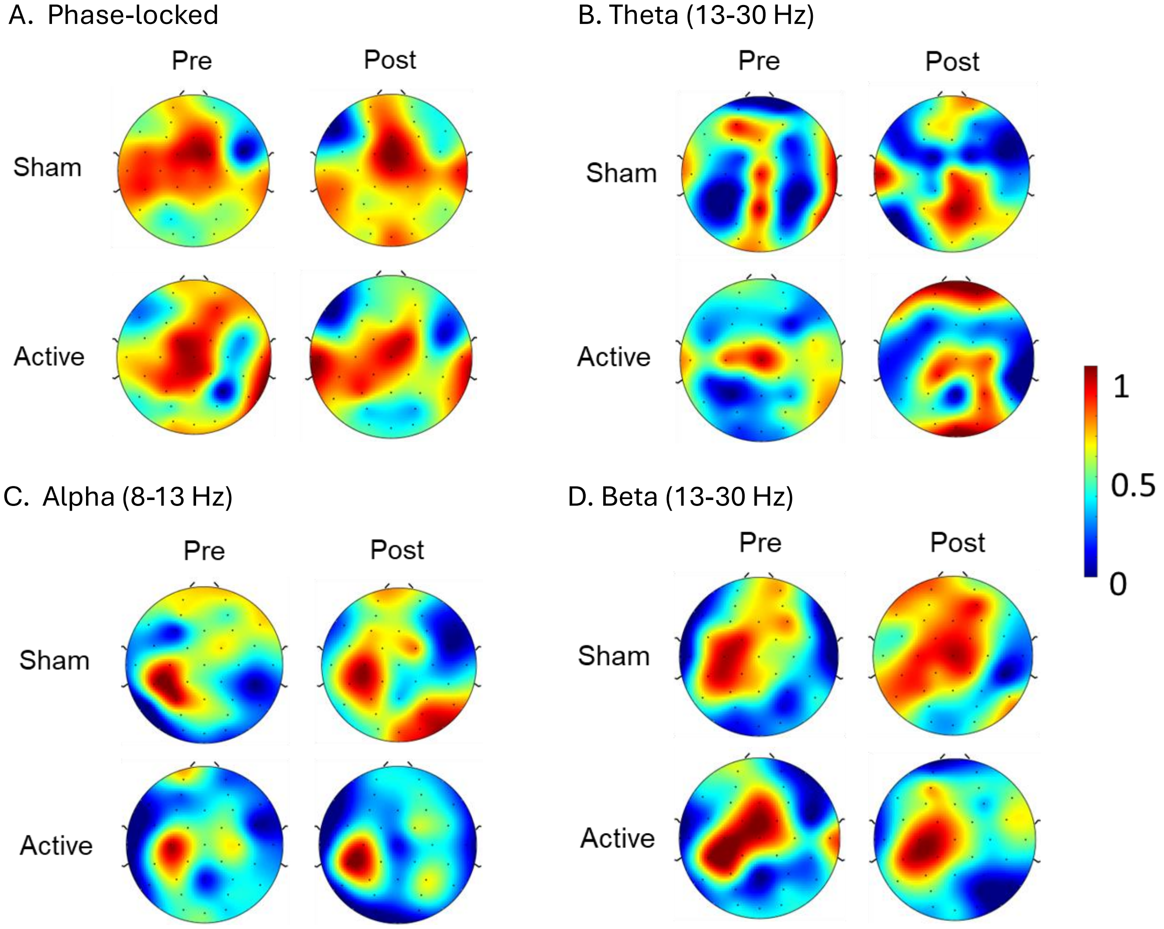


Figure S2. Topographical maps showing the strongest phase-locked and ongoing oscillatory modulations in response to periodic thermonociceptive stimulation for each condition and time point. A. For the phase-locked response the significant modulations were found at Fz (pre-sham), FC1 (post-sham), Cz (pre-active and post-active). B. For the theta band the most significant modulations were found at Pz (pre-sham and post-sham), Cz (pre-active), and Fp2 (post-active). C. For the alpha band the most significant modulations were found at P3 (pre-sham), CP5 (post-sham), C3 (pre-active), and CP5 (post-active). D. Finally, for the beta band the most significant modulations were found at C3 (pre-sham, post-sham, and post-active), and FC1 (pre-active). Color scales represent test statistic.

Table S1. Summary of Linear Mixed Model (LMM) results for all participants (n = 38), females (n = 19), and males (n = 19).

| | All (n=38) | | | | Female (n=19) | | | | Male (n=19) | | | |
|---|---|---|---|---|---|---|---|---|---|---|---|---|
| | WDT | HPT | Intensity | Painfulness | WDT | HPT | Intensity | Painfulness | WDT | HPT | Intensity | Painfulness |
| Condition | F=0.0, p=0.93 | F=0.1, p=0.73 | F=4.9, p=0.03 | F=0.21, p=0.64 | F=2.4, p=0.12 | F=1.6, p=0.21 | F=0.1, p=0.73 | F=4.5, p=0.04 | F=2.0, p=0.15 | F=1.4, p=0.23 | F=6.7, p=0.01 | F=1.4, p=0.22 |
| Time | F=9.5, p<0.01 | F=11.7, p<0.001 | F=22.7, p<0.0001 | F=18.0, p<0.0001 | **F=10.3, p<0.01** | **F=17.0, p<0.0001** | F=12.9, p<0.001 | F=13.8, p<0.001 | F=1.9, p=0.16 | F=0.9, p=0.32 | F=11.2, p<0.01 | F=8.2, p<0.01 |
| Interaction | F=0.9, p= 0.35 | F=0.1, p=0.78 | F=0.31, p=0.57 | F=0.25, p=0.61 | F=2.7, p=0.10 | F=1.0, p=0.31 | F=0.1, p=0.80 | F=0.8, p=0.37 | F=0.0, p=0.87 | F=1.4, p=0.22 | F=0.8, p=0.37 | F=0.0, p=0.92 |

Table S2. Summary of Linear Mixed Model (LMM) results for participants receiving tACS at SM-IAF. Results are presented for all participants (n = 26), females (n = 14), and males (n = 12).

| | | SM-IAF (n=26) | | | | SM-IAF-Female (n=14) | | | | SM-IAF-Male (n=12) | | | |
|---|---|---|---|---|---|---|---|---|---|---|---|---|---|
| | | WDT | HPT | Intensity | Painfulness | WDT | HPT | Intensity | Painfulness | WDT | HPT | Intensity | Painfulness |
| Condition | F score | 0.3 | 0.14 | 10.2 | 1.6 | 4.1 | 3.6 | 0.5 | 8.4 | 0.8 | 0.0 | 17.7 | 1.0 |
| | p value | 0.56 | 0.71 | <0.005 | 0.2 | 0.05 | 0.06 | 0.50 | 0.005 | 0.36 | 0.98 | <0.0005 | 0.32 |
| Time | F score | 3.7 | 9.3 | 14.7 | 11.1 | 6.3 | 8.8 | 8.4 | 10.4 | 1.5 | 0.5 | 8.6 | 5.0 |
| | p value | 0.05 | 0.003 | <0.0005 | <0.005 | 0.02 | 0.004 | 0.005 | 0.002 | 0.21 | 0.46 | 0.005 | 0.03 |
| Interaction | F score | 0.1 | 0.5 | 0.8 | 0.4 | 2.2 | **3.1** | 0.2 | 2.8 | 0.1 | 0.0 | 0.9 | 0.3 |
| | p value | 0.80 | 0.47 | 0.37 | 0.51 | 0.14 | **0.08** | 0.67 | 0.10 | 0.77 | 0.83 | 0.32 | 0.61 |

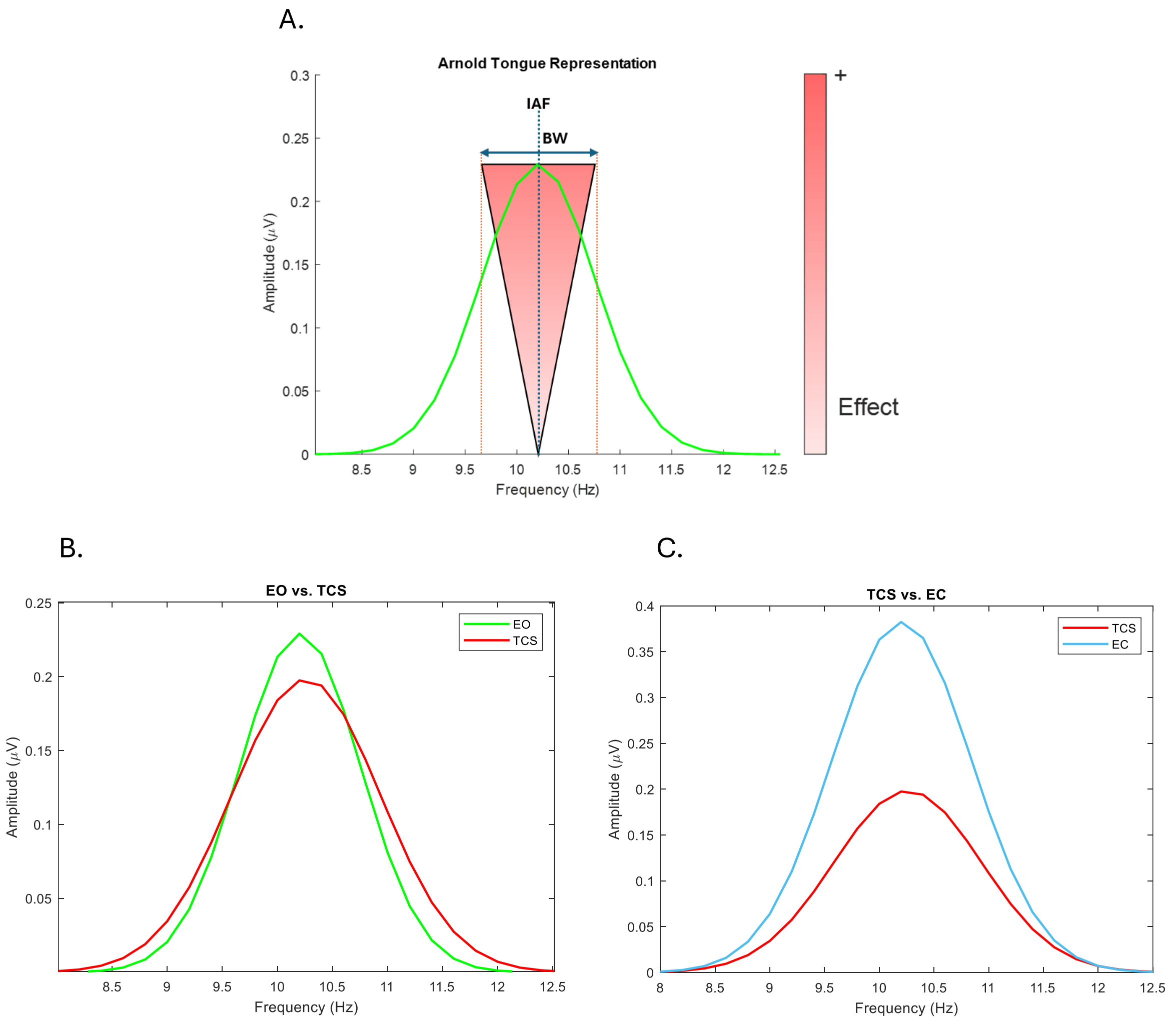


Figure S3. A. Illustration of the Arnold tongue concept. The red-shaded region represents the bandwidth (BW) of frequencies around the individual alpha frequency (IAF) that can be modulated with relative ease during transcranial stimulation. Frequencies further from the IAF require greater effort to achieve modulation. B. Comparison of BW between eyes-open (EO) and thermal cutaneous stimulation stimulation (TCS) conditions. TCS exhibits a significantly larger BW (0.25 Hz increase) compared to EO, indicating enhanced capacity for frequency modulation during TCS. This difference is statistically significant (Wilcoxon signed-rank test with Bonferroni correction, $p = 0.0007$). C. Comparison of alpha amplitudes between TCS and eyes-closed (EC) conditions. TCS shows a reduced alpha amplitude (0.17 μV lower) compared to EC. This indicates that EC requires more energy to desynchronize and pull the IAF to a new value during modulation. This difference is statistically significant (Wilcoxon signed-rank test with Bonferroni correction, $p < 0.0001$). Together, panels B and C highlight that TCS provides favorable conditions for modulating IAF, aligning with the Arnold tongue concept illustrated in panel A. Data from pre-sham condition as baseline measurements were used for this analyses. Same results can be achieved with pre-active conditions (as another baseline measurements).
